# Establishment of a Reverse Genetics System for the Study of Human Immune Functions in Mice

**DOI:** 10.1101/2025.03.26.645295

**Authors:** Priya Pal, Shuai Gao, Hongbo Gao, Marina Cella, Qiankun Wang, Liang Shan

## Abstract

Reverse genetics approaches in mice are widely utilized to understand gene functions and their aberrations in diseases. However, limitations exist in translating findings from animal models to human physiology. Humanized mice provide a powerful bridge to understanding human physiology and mechanisms of diseases pathogenesis while maintaining the feasibility of working with small animals. Methods for generating humanized mouse models that allow scientists to probe contributions of particular genes have been rudimentary. Here, we established an efficient method for generating genetically modified human cord blood derived CD34^+^ cells for transplantation, resulting in humanized mice with near-complete loss of specific gene expression by the human immune system. Mice transplanted with Cas9-edited human CD34^+^ cells recapitulate functional consequences of specific gene losses in the human immune system. This advancement enables the development of humanized mouse models with targeted gene knockouts, offering a valuable research tool for human gene function studies *in vivo*.

## Introduction

Genetically modified mice have revolutionized investigations of gene function, and regulation as well as the ability to probe into the mechanisms behind disease pathogenesis^1^. In the early 2000s, there was an increasing appreciation for the incongruencies between animal models and human physiology and disease, which launched the development of human immune system mice, also known as humanized mice. They became an invaluable tool in biomedical research, offering a unique platform for more translationally relevant work from studying underlying mechanisms of human diseases to testing potential therapeutic strategies^2,3^. There are two common ways to ‘humanize’ a mouse, use of humanized genetically engineered model whereby a human gene replaces the mouse ortholog via targeted gene replacement and can be used for the narrow investigation of that particular gene or by engrafting immunodeficient mice with human hematopoietic stem and progenitor cells (HSPCs, CD34^+^ cells),^4^ leading to the development of a functional human immune system within the mouse.^5,6^ However, the current approaches for specifically knocking out genes within the human immune system of mice have faced several challenges. Although gene knockout has been achieved in human and mouse HSPCs using electroporation of Cas9 and single guide RNA (sgRNA) ribonucleoprotein (RNP) complexes *in vitro,*^7–9^ immune reconstitution in humanized mice using genome edited human HSPCs remains challenging. A recent proof-of-concept study showed that deletion of the human CD33 coding gene in CD34^+^ cells resulted in development of CD33 deficient monocytes in mice,^8^ demonstrating the possibility of interrogating gene-specific functions within the human immune system in mice. Here, we aimed to establish an efficient method for genome editing in human CD34^+^ cells derived from umbilical cord blood and assess the impact of these modifications in a humanized mouse model. We optimized the efficiency of gene knockout in human CD34^+^ cells using electroporation of Cas9/sgRNA ribonucleoprotein (RNP) complexes. We achieved nearly 100% knockout in mice that received edited human CD34^+^ cells, generating human transgenic mice on both an NSG-SGM3^10^ and MISTRG-6-15^11,12^ backgrounds, while attaining no limitations in engraftment levels compared to controls. Furthermore, we generated *RAG2*-, *TCF7*-, *CCR5-,* and *IFNAR-*knockout humanized mice and examined the impact of these deletions on hematopoiesis and HIV-1 infection. In summary, our study offers an efficient method for genome editing in human HSPCs, highlighting its significant potential for studying gene function and antiviral immunity in humanized mouse models.

## Results

### Optimizing gene knockout efficiency in human cord blood derived CD34^+^ cells by electroporation of Cas9/sgRNA RNP complexes *in vitro*

Electroporation of Cas9/sgRNA RNP complexes is a powerful technique for genome editing in human primary cells.^13–15^ The exact parameters used for electroporation determine the ability to obtain high transduction efficiencies and more importantly high gene disruption efficiencies. We optimized various parameters, such as pulse code, as well as the concentrations and ratios of the Cas9 protein and sgRNA for effective genome editing for human CD34^+^ cells. To determine optimal gene knockout efficiency in human HSPCs, we first tested different pulse codes using the Lonza 4D electroporation system. To do this, we targeted the *CD45* coding gene, due to its trackability via flow cytometry with two sgRNAs. Cord blood HSPCs were electroporated with Cas9/sgRNA RNP complexes and subsequently cultured for five days in the presence of stem cell cytokines. Flow cytometry was employed to assess CD45 knockout efficiency and cell viability (**Figure 1A**). Among the tested pulse codes, program code DZ-100 resulted in the highest and most consistent editing efficiency (**Figures 1B** and **1C**). Specificity, this approach achieved nearly 100% CD45 knockout across various donor samples while maintaining approximately 65% cell viability (**Figures 1C and 1D**). Next, we optimized the doses of the Cas9 protein and sgRNAs to achieve maximal gene knockout. We observed that sgRNA concentrations below 25 μM in the 20 μL reaction system significantly reduced the knockout efficiency (**Figures 1E-1G**). Consequently, for our subsequent experiments, we selected the DZ-100 program and maintained Cas9 and sgRNA concentrations at 10 μM and 25 μM, respectively. Under these conditions, we ensured maximum gene knockout efficiency while preserving acceptable levels of cellular viability.

**Figure 1.**
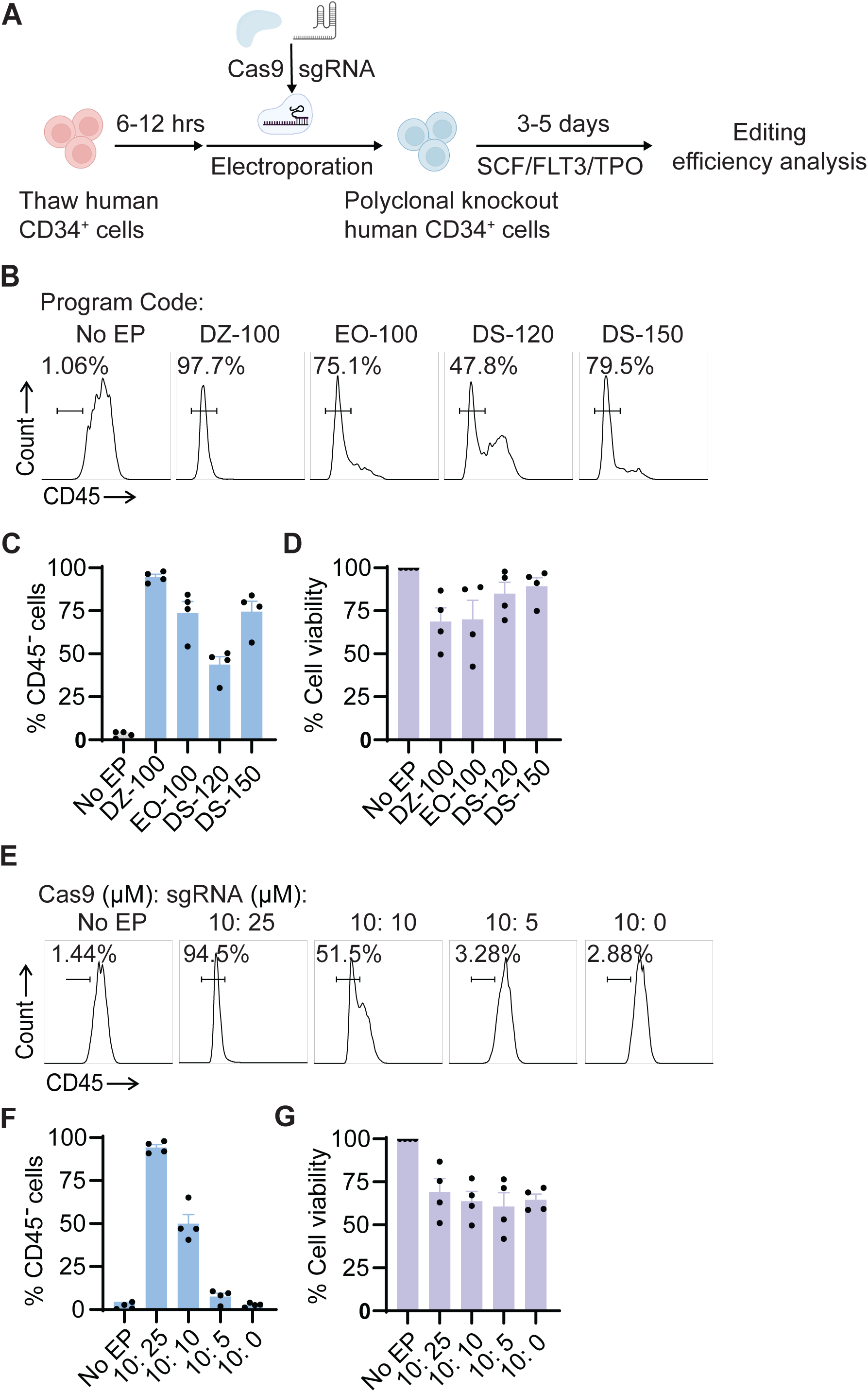
Optimized electroporation of Cas9/sgRNA RNPs in human CD34^+^ HSPCs. NSG-SGM3 mice were used in this figure. (A) Experimental scheme of CRISPR/Cas9 gene knockout in human CD34^+^ HSPCs. (B-D) Optimization of the electroporation program. Representative FACS plots of CD45 expression (B), percentage of edited (CD45^-^) cells (C), and percentage of viable cells after electroporation (D) are shown. Human CD34^+^ cells were electroporated with Cas9/sgRNA RNP complexes targeting human CD45 by using different program codes. Electroporated cells were cultured with SCF (100 ng/ml), FLT3 (100 ng/ml), and TPO (100 ng/ml) for 5 days before flow cytometry analysis. (E-G) Optimization of the input sgRNA amounts. Human CD34^+^ cells were electroporated in the presence of various amounts of Cas9/sgRNA complexes and subsequently CD45 expression and cell viability were measured as above. Data are shown as mean ± SEM of four donors (n = 4).

### Optimizing immune reconstitution by electroporated human cord blood CD34^+^ cells in immunodeficient mice

One of the most common approaches to creating humanized mice involves transferring human HSPCs into immunocompromised mice. Given the cell viability loss during electroporation, we began by investigating how the initial number of electroporated cells and the *in vitro* recovery post electroporation impacts the engraftment levels (**Figure 2A**). Specifically, we engrafted 3,000, 10,000, or 30,000 electroporated human cord blood CD34^+^ cells into newborn NSG-SGM3 mice. These cells were electroporated with the Cas9 protein and non-targeting sgRNAs. Human immune reconstitution was evaluated 12 to 14 weeks post-engraftment by analyzing the presence of human CD45^+^ (huCD45^+^) cells in peripheral blood. Our findings indicate a direct correlation between a higher initial cell count and improved engraftment rates in the blood, spleen, and bone marrow (BM) (**Figure 2B**). We subsequently assessed whether *in vitro* culture of electroporated CD34^+^ cells enhances engraftment efficiency. Electroporated cells were cultured for durations of 0.5, 24 or 72 hours before engraftment. We observed that the percentage of human CD45^+^ cells in the spleen, bone marrow, liver, and lung exhibited no significant difference, indicating CD34^+^ HSPCs short-term *in vitro* culture did not have a substantial impact on the engraftment efficiency (**Figure 2C and D**). Moreover, the proportion of human T cells, B cells, monocytes, and NK cells in tissues was not impacted by the duration of *in vitro* culture (**Figure 2E and F**). Consequently, we opted to transplant a minimum of 30,000 electroporated cells with a 3-day period of in vitro culture in our subsequent experiments. The 3-day in vitro culture period provided the flexibility needed to wait for the delivery of newborn mice.

**Figure 2.**
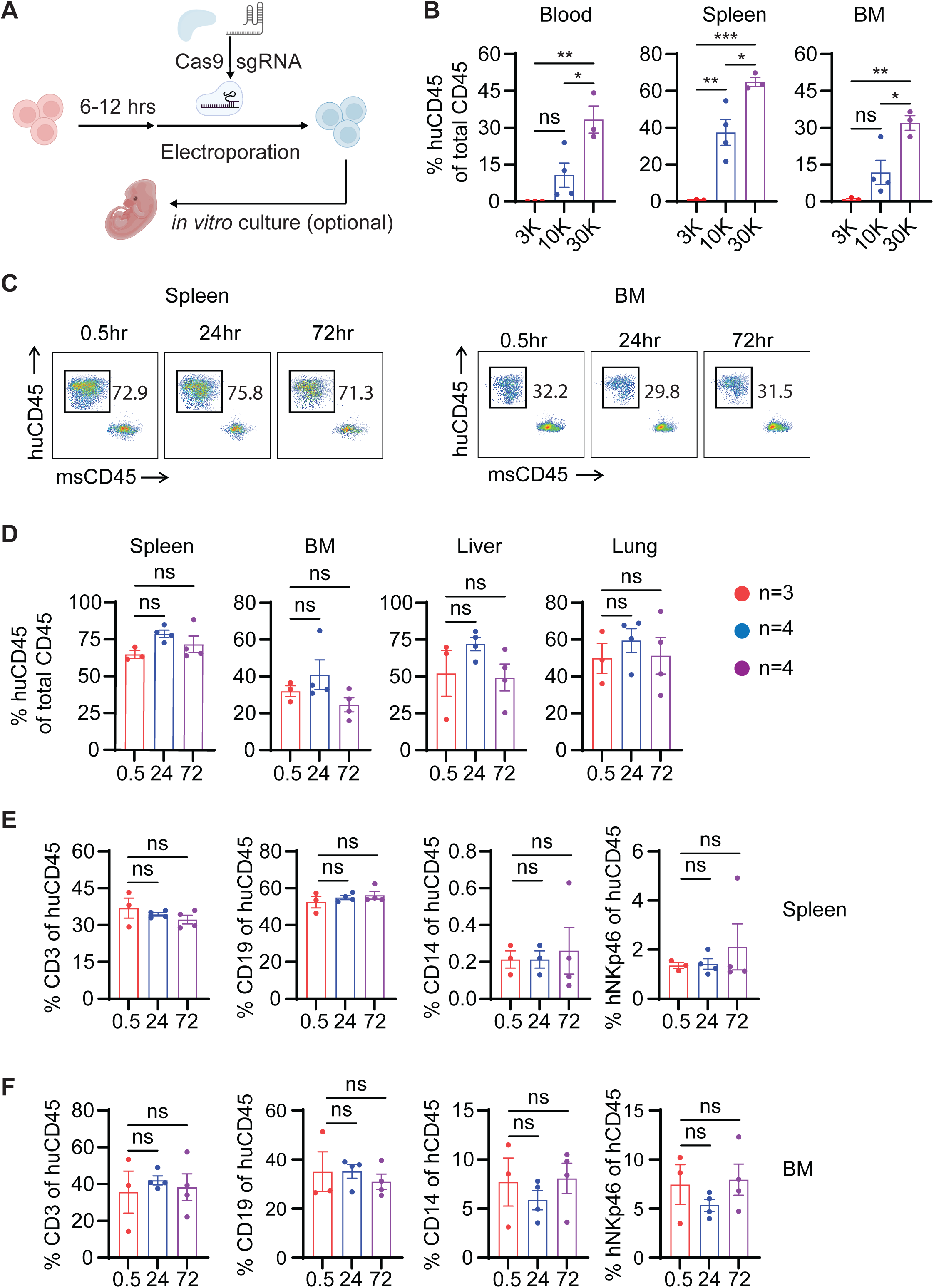
Optimized cell numbers for immune reconstitution and multilineage differentiation after transplantation of electroporated human CD34^+^ cells into immunodeficient mice. NSG-SGM3 mice were used in this figure. Samples were harvested 14 weeks post engraftment. (A) Experimental scheme of electroporation and transplantation procedures. (B) Percentages of human CD45^+^ (huCD45^+^) cells in the blood, spleen, and BM of indicated groups. Data are presented as the mean ± SEM. 3K (n = 3), 10K (n = 4), and 30K (n = 3). p values were calculated using the one-way ANOVA with Tukey’s multiple comparisons tests. * p < 0.05, ** p < 0.01, and ***p < 0.001. (C-F) Human immune cell engraftment and multilineage differentiation using *in vitro* cultured CD34^+^ cells post electroporation. After electroporation, CD34^+^ cells were cultured *in vitro* for 0.5, 24 or 72 hours with the presence of SCF (100 ng/ml), FLT3 (100 ng/ml), and TPO (100 ng/ml) before engraftment. Tissues were harvested 14 weeks after engraftment. Reconstitution of total human immune cells (C and D) and multilineage differentiation (E and F) were determined by FACS. Data are presented as the mean ± SEM. *p* values were calculated using the one-way ANOVA with Tukey’s multiple comparisons tests.

### Immune reconstitution by Cas9-edited human cord blood CD34^+^ cells in immunodeficient mice

We next aimed to evaluate human immune reconstitution after optimizing the electroporation of Cas9/sgRNA RNP complexes and engraftment conditions. To this end, we targeted the *HLA- A2* gene, which is not involved in hematopoiesis and can be easily detected via flow cytometry. We first validated the editing efficiency *in vitro* and found that nearly 100% knockout efficiency of HLA-A2 in CD34^+^ cells was achieved five days after Cas9/sgRNA RNP electroporation (**Figure S1A**). Subsequently, we transplanted these *HLA-A2* edited and Cas9 control (Cas9 ctrl) cells into newborn NSG-SGM3 mice and assessed the *HLA-A2* knockout efficiency 14 weeks post engraftment. We observed a complete loss of human HLA-A2 expression in the blood, spleen, BM, liver, and lung of each mouse engrafted with the edited cells (**Figures 3A and B**). In addition, we observed that the levels of reconstitution of human CD45^+^ cells in the blood, spleen, BM, liver, and lung of engrafted mice were comparable between the Cas9 control and *HLA-A2* targeted groups, suggesting that the knockout of *HLA- A2* had no significant impact on the stemness and differentiation potential of edited CD34^+^ cells (**Figures 3C and D**, **and S1B**). Hence, these results illustrate that our methodology successfully led to the creation of transgenic humanized mice with selective human gene knockout without impairment in overall engraftment.

**Figure 3.**
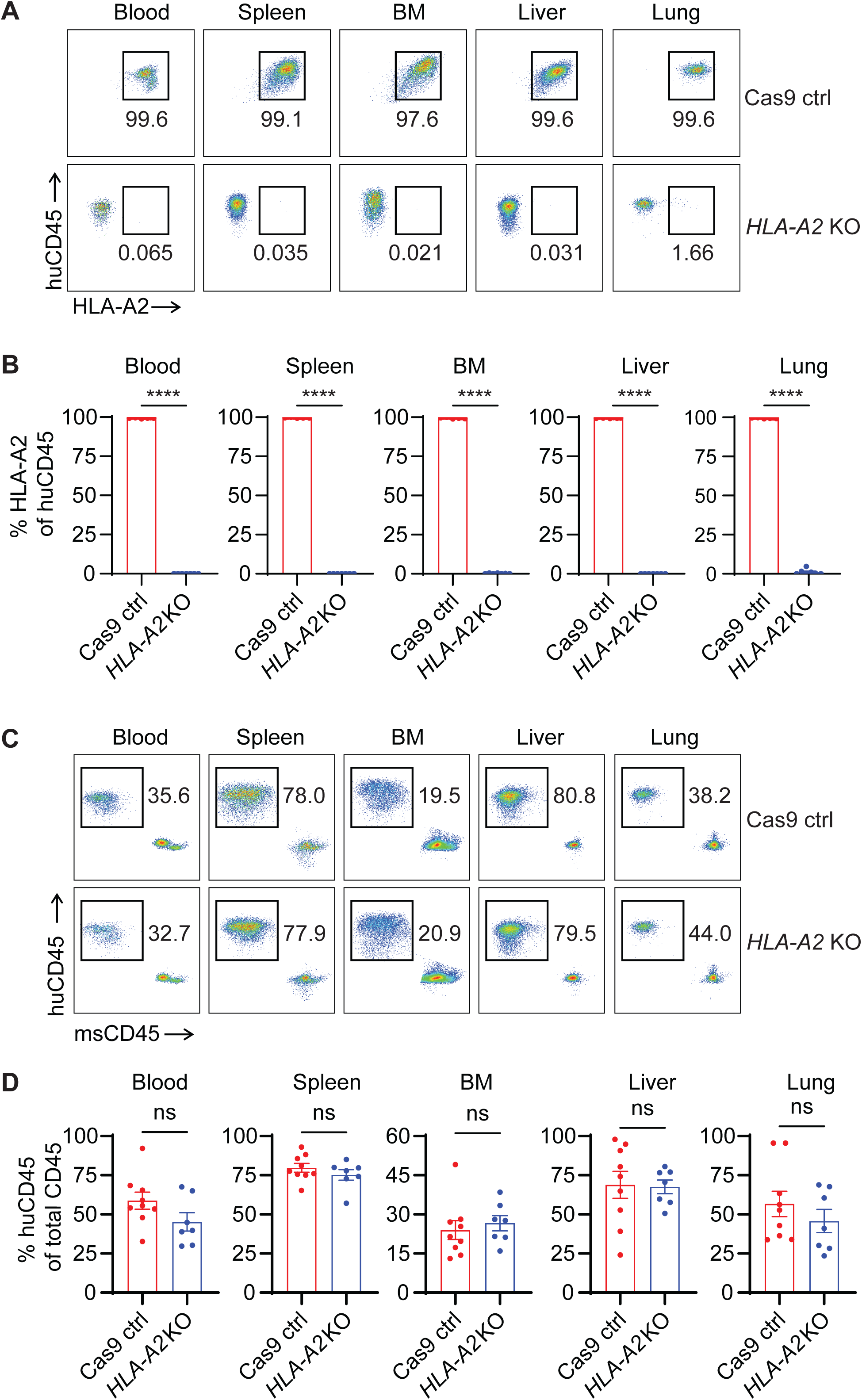
Immune reconstitution by *HLA-A2*-KO CD34^+^ cells in immunodeficient mice. NSG-SGM3 mice were used in this figure. Samples were harvested 14 weeks post engraftment. (A) Representative FACS plots of human HLA-A2 expression by huCD45^+^ cells in mice 14 weeks after engraftment in blood, spleen, BM, liver, and lung. (B) Percentages of human HLA-A2^+^ of huCD45^+^ cells in the blood, spleen, BM, liver, and lung. (C) Representative FACS plots of human (huCD45^+^) and mouse (msCD45^+^) immune cells in the blood, spleen, BM, liver, and lung. (D) Human immune cell reconstitution in the blood, spleen, BM, liver, and lung. Data were shown as the mean ± SEM. Cas9 ctrl (n = 9), *HLA-A2* KO (n = 7). *p* values were calculated using an unpaired two-tailed t-test. ****p < 0.0001.

Next, we aimed to evaluate whether CRISPR-Cas9 edited HSPCs provided a reverse genetics system which faithfully recapitulated known immune defects when essential genes were deleted. To do this, we used the MISTRG-6-15 mouse strain, which supports the development of human innate immune system more efficiently than the NSG-SGM3 mouse strain, so that the loss of T cells and B cells caused by targeted gene deletions would not result in a complete loss of human immune reconstitution. In mice, loss of T cell factor 1 (Tcf1, encoded by *Tcf7*) blocks thymocyte differentiation but has no direct impact on humoral responses.^16,17^ We introduced *TCF7*-KO and Cas9 ctrl cord blood derived CD34^+^ cells into MISTRG-6-15 mice to evaluate immune reconstitution. In mice engrafted with *TCF7*-KO CD34^+^ cells, a complete loss of T cell development was observed while B cells, myeloid cells, and NK cells remained unaffected (**Figures 4A-C**). A slight increase of B cell or NK cell frequency in the *TCF7*-KO group was likely a reflection of the loss of T cells. We confirmed that targeted deletion of *TCF7* had no impact on hematopoiesis, as evidenced by the comparable levels of bone marrow progenitors between the *TCF7*-KO and Cas9 control groups including total CD34^+^ cell as well as different subsets of progenitor cells (**Figures 4D and E**). These results indicate a specific ablation of mature T cells by *TCF7*-KO. Next, we introduced *RAG2*-KO and Cas9 ctrl cord blood derived CD34^+^ cells into MISTRG-6-15 mice to evaluate the reconstitution of adaptive immunity (**Figures 4F**). Both B cell and T cell populations were absent in mice engrafted with *RAG2*-KO CD34^+^ cells, confirming the essential role of RAG2 in genomic rearrangement of B cell and T cell receptors.^18^ Notably, the *RAG2*-KO animals are excellent tools to study human innate immune responses to pathogens or other diseases. Our *TCF7*-KO and *RAG2*-KO transgenic humanized mice demonstrate that CRISPR -Cas9 edited human HSPCs provide a robust reverse genetics system which will efficiently allow scientists to probe the function and contribution of a variety of immune genes *in vivo* in the broader context of a complete immune system.

**Figure 4.**
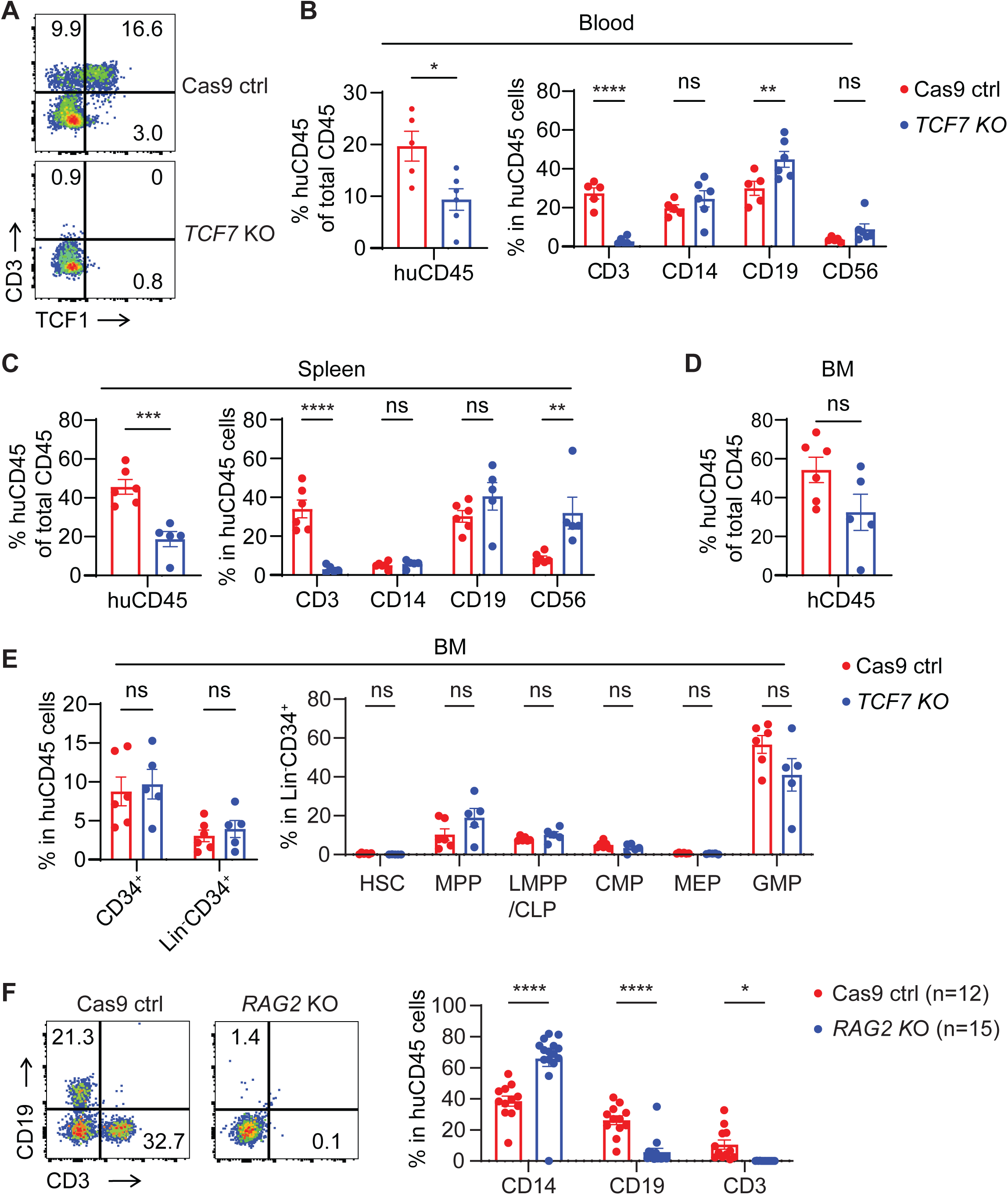
The human immune systems developed from *TCF7*-KO and *RAG2*-KO CD34^+^ cells recapitulate established immune deficiencies. MISTRG-6-15 mice were used in this figure. Samples were harvested 12 weeks post engraftment. (A) Representative FACS plots of TCF1 expression by huCD45^+^ cells. (B-D) Reconstitution of human immune cells in blood (B), spleen (C) and bone marrow (D) of mice engrafted with *TCF7*-KO and Cas9 ctrl CD34^+^ cells. (E) Analysis of bone marrow total huCD45^+^ cells and human hematopoietic stem and progenitor cells from mice engrafted with *TCF7*-KO and Cas9 ctrl CD34^+^ cells. (F) T cell, B cell, and myeloid cell development in mice engrafted with *RAG2*-KO and Cas9 ctrl CD34^+^ cells. Representative FACS plots of CD3 and CD19 expression by blood huCD45^+^ cells were shown. Frequency of CD14^+^, CD19^+^ and CD3^+^ blood cells were determined by FACS.

### Mice with an edited human immune system have altered responses to HIV-1 infection

The humanized mouse reverse genetics system may offer new insights into HIV-1 and host interactions *in vivo*. CCR5 and CXCR4 are the two co-receptors for HIV entry,^19,20^ and CCR5- tropic HIV-1 isolates are transmitted by all routes and almost always dominate throughout the course of infection prior to the onset of AIDS.^21,22^ We introduced human *CCR5*-KO and Cas9 ctrl CD34^+^ cells into NSG-SGM3 mice. The successful establishment of the *CCR5* knockout human immune system in the mice was confirmed using flow cytometry and deep sequencing analysis. These methods revealed that more than 95% of human cells were edited at the *CCR5* locus while total human cell engraftment remained unaffected (**Figures 5A-C** and **S2A**). We then infected these mice with the CCR5-tropic HIV_BaL_ and monitored the levels of plasma HIV- 1 RNA and the CD4^+^ T-cell loss in the blood. Mice that were reconstituted with the *CCR5*-KO immune system displayed undetectable levels of plasma HIV-1 RNA (**Figures 5D**), and CD4^+^ T-cell loss was prevented in these animals (**Figure 5E**). Moreover, both the frequency and the number of CD4^+^ T cells in tissues including spleen, BM, liver, and lung were higher in mice with a *CCR5*-KO immune system in comparison to the Cas9 control mice (**Figures 5F and G**). Type I interferons (IFN) play a complicated role in HIV-1 infection, as they not only suppress viral replication, but also drive immune activation and exhaustion.^23^ On one hand, IFN-α monotherapy during clinically asymptomatic HIV-1 infection without antiretroviral therapy reduced plasma viral loads, prevented CD4 decline, and delayed disease progression to AIDS.^24–26^ On the other hand, *in vivo* blockade of IFN-α receptor (IFNAR) in SIV-infected rhesus macaques dampened upregulation of IFN-stimulated genes, which led to heightened plasma viremia and accelerated CD4 decline.^27^ Although these interventions demonstrate strong anti-HIV-1 activities of type 1 IFNs, the role of type 1 IFNs induced during natural HIV- 1 infection *in vivo* remains unclear. We introduced human *IFNAR*-KO and Cas9 ctrl CD34^+^ cells into MISTRG-6-15 mice, as this mouse strain develops myeloid cells more efficiently than NSG-SGM3. We confirmed nearly 100% editing efficiency of *IFNAR* (**Figure 6A and S2B**) and observed that the levels of reconstitution of human CD45^+^ cells as well as T cell development were comparable between the Cas9 control and *IFNAR* KO groups (**Figure 6B**-**D**). Upon HIV-1 infection, both groups of mice produced detectable and comparable levels of IFN-α2 (**Figure 6E**). Importantly, the plasma HIV-1 RNA levels were 10 to 100-fold higher in the *IFNAR* KO group (**Figure 6F**), which suggests that type 1 IFNs contribute substantially during acute HIV-1 infection to control viral replication.

**Figure 5.**
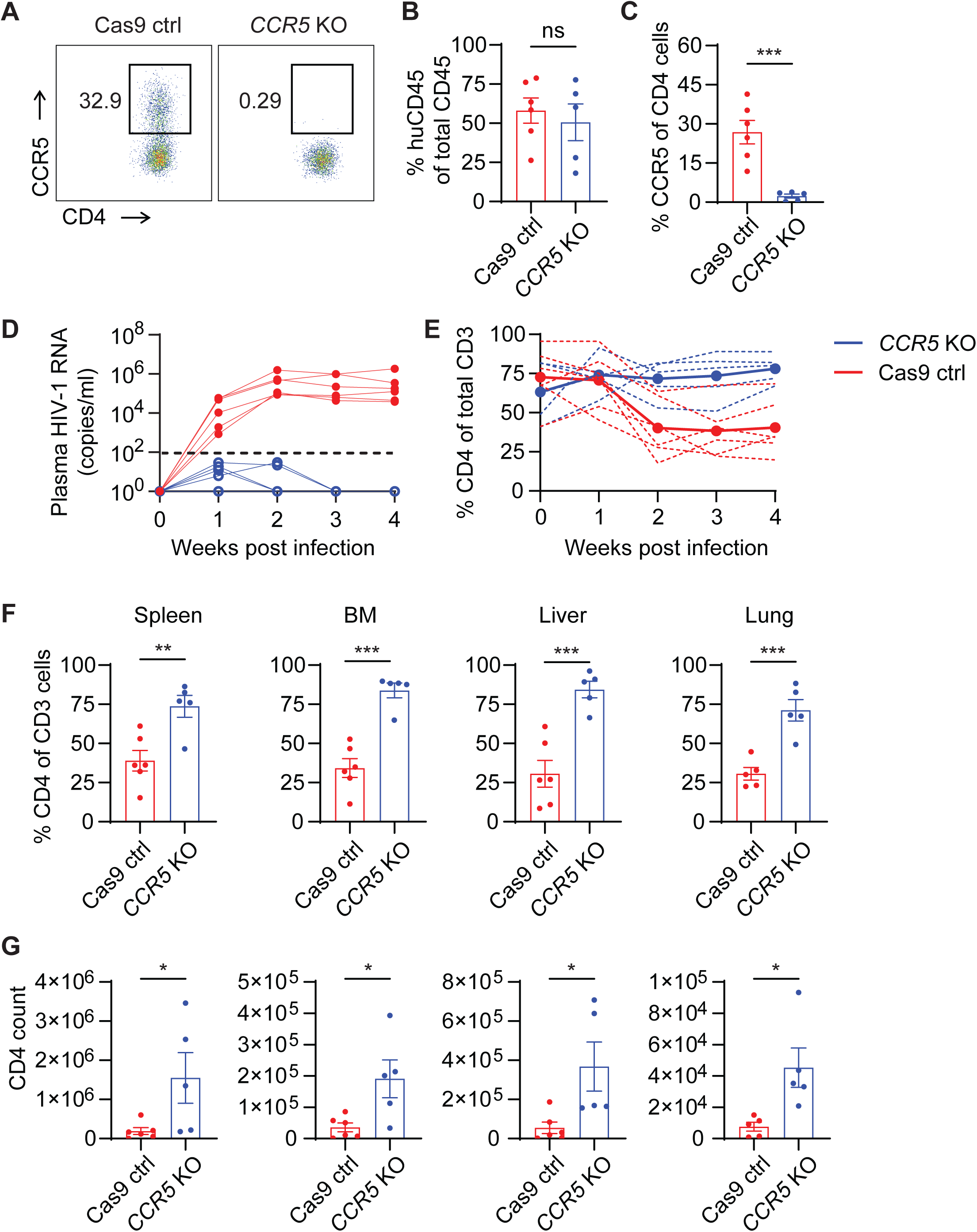
Mice with a human immune system developed from *CCR5*-KO CD34^+^ cells are resistant to HIV-1 infection. NSG-SGM3 mice were used in this figure. Mice were infected with HIV_BaL_ 14 weeks post engraftment. (A) Representative FACS plots of CCR5 expression in blood CD4^+^ T cells before infection. (B-C) Percentage of huCD45^+^ cells (B) and CCR5^+^ CD4 T cells (C) in blood before infection. (D) Plasma viral loads measured by RT-qPCR. (E) Loss of blood CD4^+^ T cells determined by FACS. (F-G) The frequency and number of human CD4^+^ T cells in tissues. Data were shown as the mean ± SEM. Cas9 ctrl (n = 6), *CCR5* KO (n = 5). *p* values were calculated using an unpaired two-tailed t-test. * p < 0.05, ** p < 0.01, and ***p < 0.001

**Figure 6.**
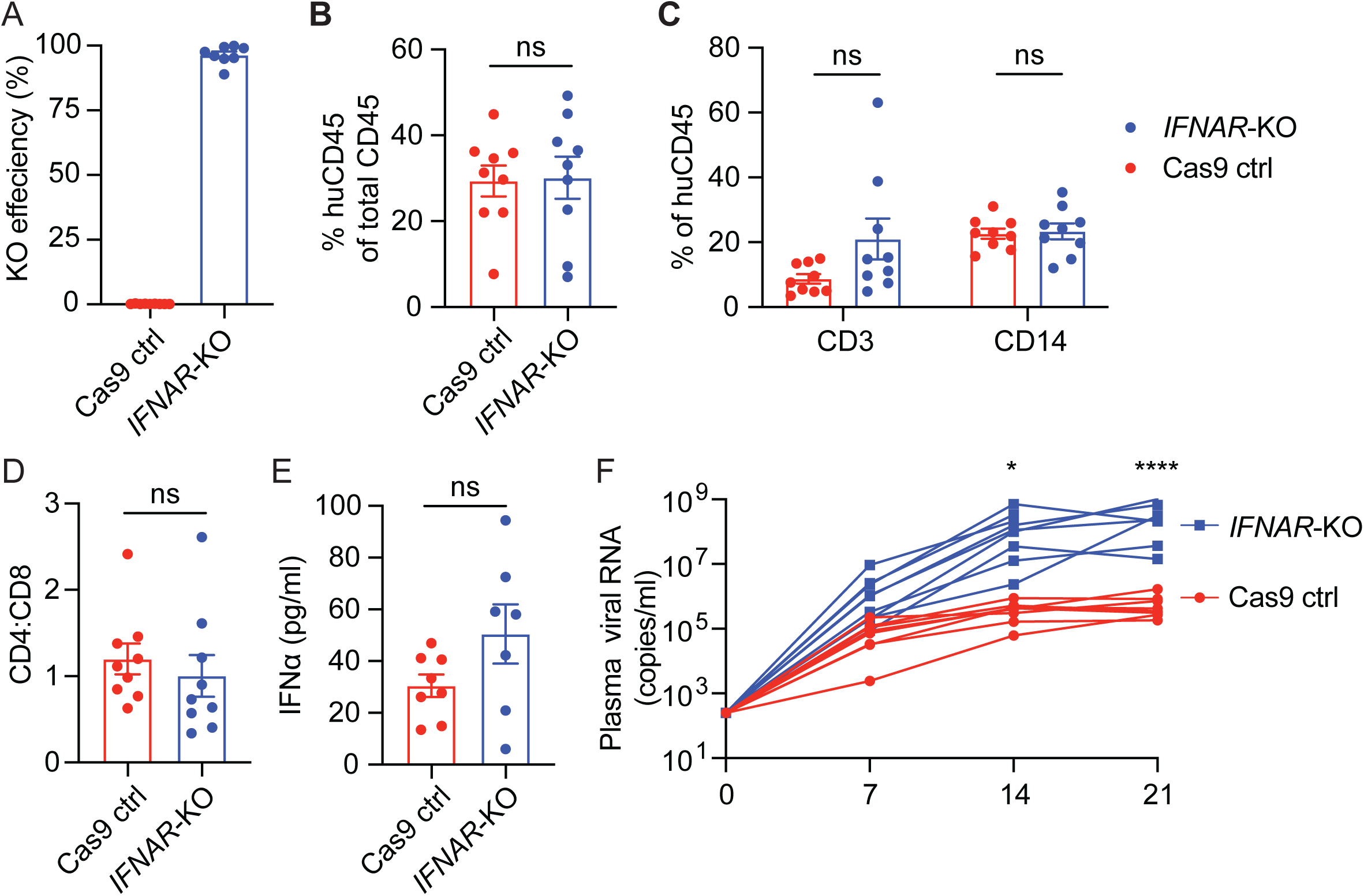
Type I interferons suppress HIV-1 replication *in vivo*. MISTRG-6-15 mice were used in this figure. Mice were infected with HIV_BaL_ 12 weeks post engraftment. (A) Editing efficiency. PBMCs were collected for genomic DNA extraction. Editing efficiency was determined by the MiSeq Illumina sequencing of the targeted region (see also Fig. S2B). (B-D) Percentage of huCD45^+^ (B), CD3^+^ and CD14^+^ (C), and CD4:CD8 ratio (D) cells in the blood in the indicated groups. (E) Plasma IFNα2 levels in infected and control mice measured by ELISA. (F) Plasma viral loads were measured from 1 week to 3 weeks after infection by RT-qPCR.

## Discussion

Here, we described a reverse genetics system which allows us to interrogate specific gene functions in the development of human immune system and its contribution to controlling human pathogens. We used well-characterized immune genes as a proof-of-concept to demonstrate highly efficient genome editing and immune reconstitution of transgenic HSPCs in mice which provides a successful reverse-genetics model. We chose the MISTRG-6-15 and the NSG-SGM3 mice as they support human hematopoiesis efficiently due to their production of multiple human cytokines and growth factors. While NSG mice are the most commonly used strain for the generation of humanized mice, 50,000 to 200,000 unedited CD34^+^ cells per mouse are needed for human immune reconstitution in the NSG model, which would be prohibitive when using transgenic cells.

Genetical ablation of certain human genes in primary CD4^+^ T cells to study HIV-1 and host interaction can provide valuable insights into HIV-1 biology that help identify novel therapeutic targets.^28,29^ In comparison with the functional studies *in vitro*, the platform described here has notable advantages. For example, immune cells residing in different tissues often have distinct phenotypes and functions because they interact with tissue-specific parenchymal cells and rely on different cytokines for maturation and survival.^30–36^ Although latent HIV-1 can be found in tissue macrophages,^37–40^ without an animal model lacking a T cell reservoir, it remains unclear whether tissue macrophages contribute to HIV-1 reservoirs and whether there would be viral rebound after analytic treatment interruptions. The *RAG2*-KO and *TCF7*-KO models we describe can serve as T-cell deficient humanized mice to understand the role of tissue macrophages in HIV-1 persistence. Additionally, disruption of certain immune functions such as T cells or type I IFN response could help understand how the human immune system controls pathogens and how pathogens evolve under immune selective pressure.

Notably, there are still limitations with regard to broad applications of the reverse genetics in humanized mice. First, genome editing cannot be used to study the function of genes that are essential for hematopoiesis. Second, unlike the commonly used conditional knockout systems such as the Cre recombinase, the current genome editing of the human immune system applies to all immune cells without cell type or tissue specificity. Furthermore, since the number of human CD34^+^ cells from a standard cord blood sample is limited, multi-arm large cohort studies remain challenging. Nonetheless, this study represents a significant step forward in the development of a robust platform to functionally study the contribution of various genes to the control of HIV pathogenesis and to directly probe the host-pathogen interface using humanized mice.

## Materials and Methods

### Mice Strain

NSG-SGM3 mice were purchased from Jackson Laboratory (Strain #013062) and the colony was maintained at Washington University School of Medicine. The MISTRG-6-15 human cytokine knock-in mice was described previously^12^. The mouse colony was maintained at Washington University School of Medicine. Both male and female mice were included. All animal experiments were approved by the Institutional Animal Care and Use Committee of Washington University School of Medicine.

### Human samples

Anonymous human cord blood samples were collected at the Cleveland Cord Blood Center.

### Plasmids and viruses

The replication-competent HIV_BaL_ was produced by infecting CD8-depleted PHA-stimulated PBMCs. The culture supernatant was collected six to nine days post-infection. NL4-3-BAL was generated by replacing the BAL-01 envelope into the NL4-3 consensus sequence and was used for infections of *IFNAR*-KO mice. Virus was generated by transfection of 293T cells with plasmids and purified and aliquoted using LentiX at -80C. Virus was quantified using p24 ELISA kit and 10ng/mouse was used for infection.

### CRISPR knockout in CD34^+^ cells

EasySep™ Human Cord Blood CD34 Positive Selection Kit III (Stem Cell Technologies # 17897) was used to purify CD34^+^ cells from cord blood. Cas9-sgRNA ribonucleoprotein (RNP) complexes were electroporated into CD34^+^ cells using the Lonza 4D Nucleofector system. The recombinant Cas9 protein was obtained from IDT. The modified synthetic sgRNAs were purchased from Synthego and the sequences are listed in **Table S1**. RNP complexes were prepared by mixing sgRNA (100 pmol) with Cas9 (40 pmol) and incubating them for 10 minutes at room temperature. 0.2 × 10^6^ CD34^+^ cells were washed with PBS and resuspended in 20 µl buffer P3 (Lonza #V4XP-3032). To deliver two specific sgRNAs targeting one gene, 2 µl of each RNP complex were then mixed with the cell suspension and transferred into a 16- well reaction cuvette of the 4D-Nucleofector System. The CD34^+^ cells were electroporated using the programs DZ-100, unless indicated. After electroporation, CD34^+^ cells were resuspended in 100 µl of prewarmed IMDM and transferred to a 96-well plate to recover for 30 minutes at 37°C. The electroporated CD34^+^ cells were then washed with PBS before in vitro culture or transplantation into the mice.

### Flow cytometry analysis

The mouse tissue cell suspensions were first incubated with Zombie NIR in PBS for 30 minutes at 4°C, followed by incubation with Human TruStain FcX and TruStain FcX (anti-mouse CD16/32) blocking antibodies for 10 min at room temperature. The cell suspensions were then incubated at 4°C with fluorescence-conjugated antibodies for 30 minutes to stain surface antigens. In all experiments, stained cells were acquired on a BD LSR Fortessa, X20, or Accuri C6 (BD Biosciences), and data were analyzed by the FlowJo software.

For HSPC analysis, a cocktail of lineage markers including CD3, CD14, CD19, and CD56 were used. Progenitor cells were defined according to a previous study^41^.

### Generation and HIV-1 infection of humanized mice

One- to three-day-old newborn NSG-SGM3 mice were preconditioned with sublethal irradiation (100 cGy). No preconditioning was performed for MISTRG-6-15 mice. Mice then received 3-4×10^4^ unmodified or Cas9-edited CD34^+^ cells by an intrahepatic injection. Reconstitution of human CD45^+^ cells was assessed 9 to 12 weeks after engraftment. Experimental groups were assigned randomly.

Some engrafted mice were infected with HIV_BaL_ (10 ng p24/mouse) through retro-orbital injection. Blood samples were collected by retro-orbital or submandibular vein bleeding to quantify plasma HIV-1 RNA. The plasma viral RNA was extracted using a Quick-RNA Viral Kit (Zymo Research) and reverse transcribed using the ProtoScript II Reverse Transcriptase (NEB). A ten-fold serial dilution of HIV genomic DNA served as a standard for measuring plasma viral RNA by the HIV gag-based qPCR assay.

### Amplicon deep sequencing

In the first step, corresponding primers were used for the gene locus. In a second round of PCR using primers containing sample-specific barcodes and adapters, amplicons were sequenced with 2 × 150 paired-end reads with MiSeq Sequencing (Illumina). The CRISPResso software was used to analyze the deep sequencing data^42^.

## Statistical Analysis

The statistical analysis was conducted using Prism 9. Phylogenetic statistics and analysis were performed as above. The methods for statistical analysis are described in the figure legends. The error bars indicate the standard error of the mean.

## Data and code availability

All data are available in the main text or the supplementary materials.

## Acknowledgments

We thank the Regeneron Pharmaceuticals and the Richard Flavell Laboratory at Yale University for generating the MISTRG-6-15 human cytokine knock-in mice. This work was supported by NIH grants R01AI162203, R01AI155162, and R01AI176594 to LS, and in part by the CRISPR for Cure Martin Delaney Collaboratory for HIV cure UM1AI 164568, co-funded by NIAID, NIMH, NIDA, NINDS, NIDDK, and NHLB.

## Author contributions

P.P., Q.W., and L.S. designed the study and analyzed the data. P.P., S. G., H. G., M.C., and Q.W. performed animal studies. P.P., Q.W., and L.S. wrote the manuscript.

---

**Figure S1.**
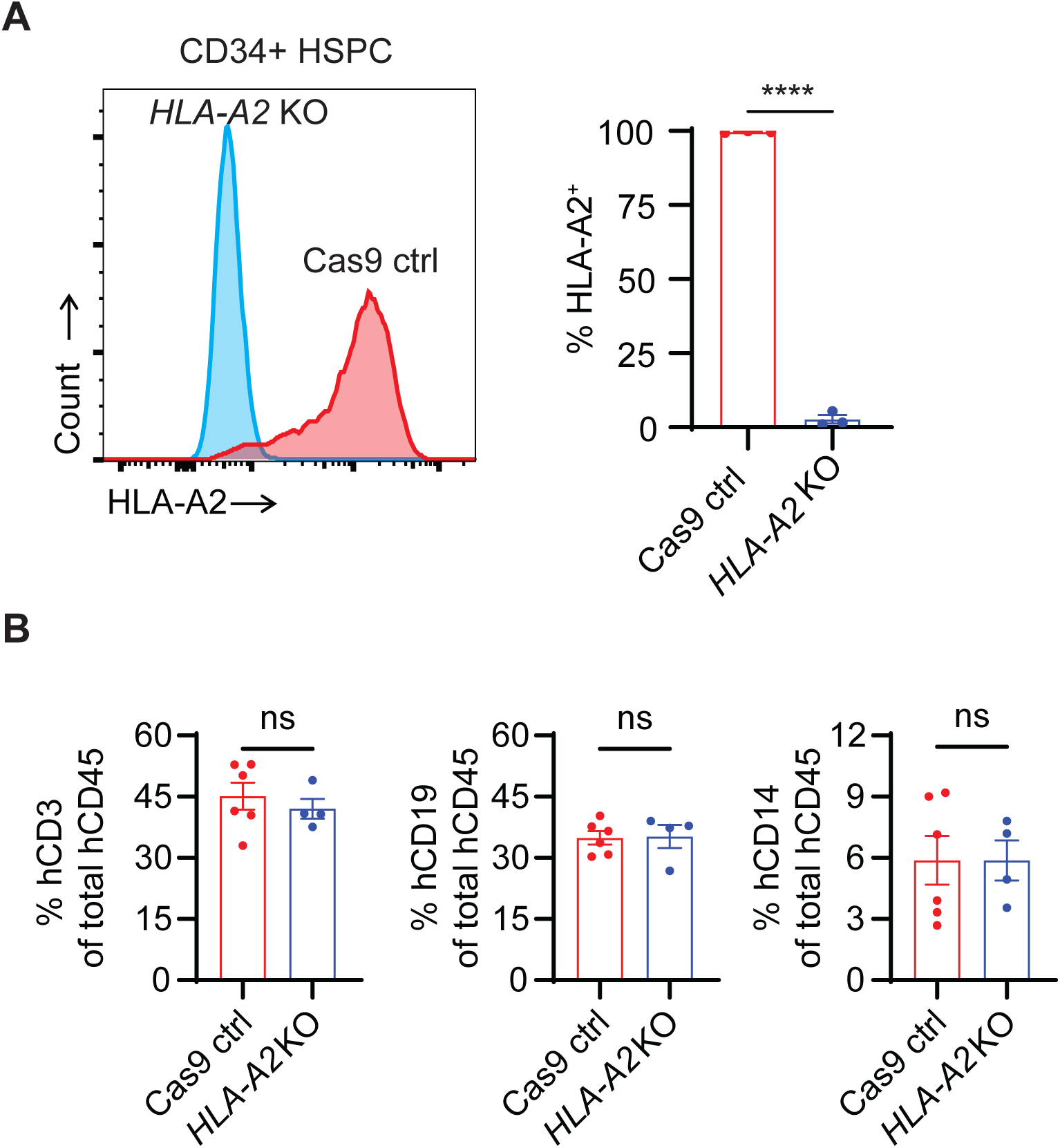
Validation of *HLA-A2*-KO CD34^+^ cells *in vitro* and in mice. NSG-SGM3 mice were used in this figure. Samples were harvested 12 weeks post engraftment. (A) Representative FACS plots and summary of human HLA-A2 expression by CD34^+^ cells. Human CD34^+^ cells were electroporated with Cas9/sgRNA RNP complexes targeting *HLA-A2*. Electroporated cells were cultured with SCF (100 ng/ml), FLT3 (100 ng/ml), and TPO (100 ng/ml) for 5 days before flow cytometry analysis. (B) Blood T cell, B cell, and myeloid cell development in mice engrafted with *HLA-A2*-KO and Cas9 ctrl CD34^+^ cells. Frequency of CD14^+^, CD19^+^ and CD3^+^ blood cells were determined by FACS. *p* values were calculated using an unpaired two-tailed t-test.

**Figure S2.**
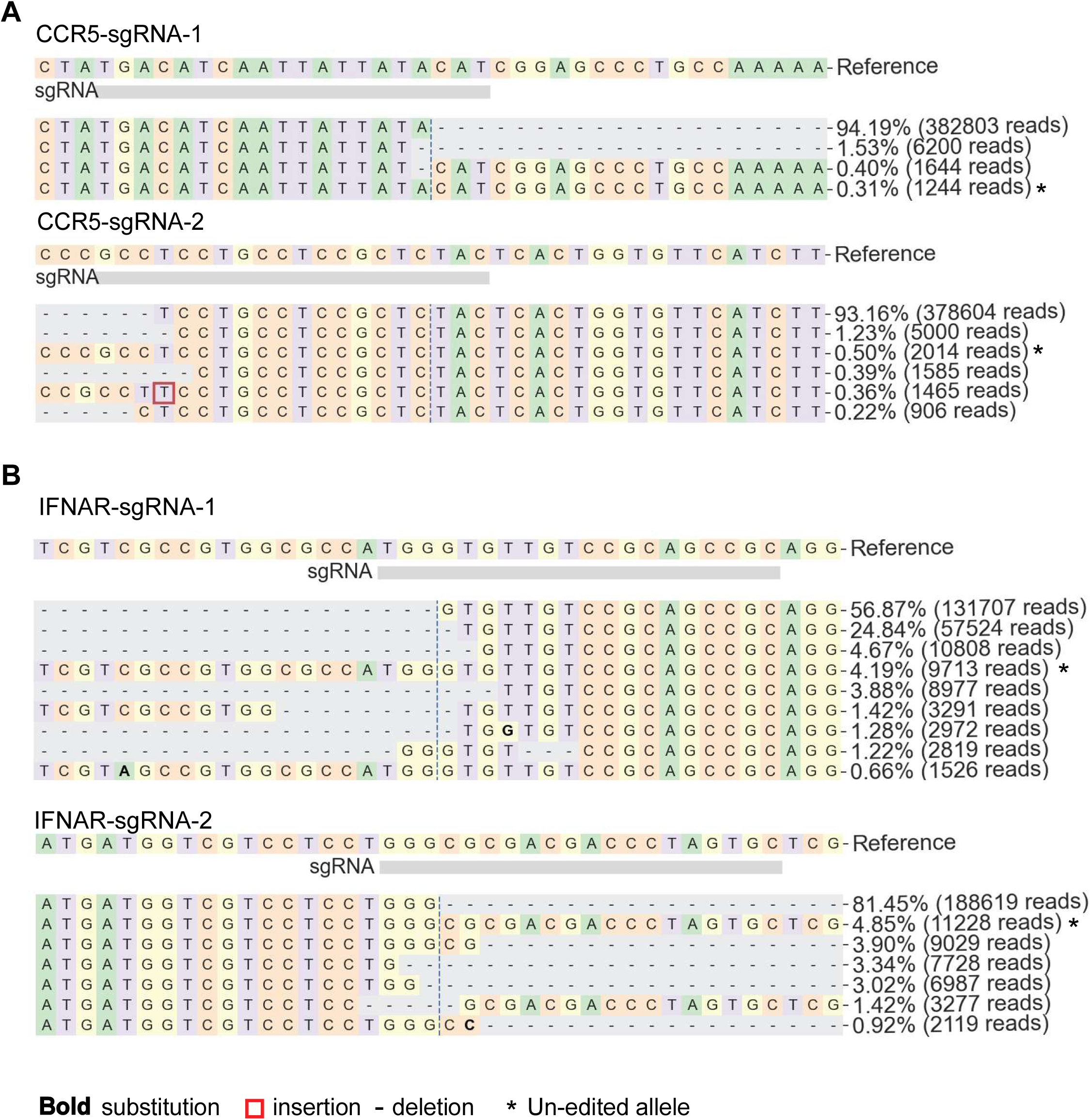
Validation of Cas9-edited CD34^+^ cells in mice by sequencing of the targeted regions. Representative alignment of the targeted regions of *CCR5* (A) and *IFNAR* (B). Library preparation and sequencing were performed using PBMCs collected 9 weeks post engraftment.

**Table S1.**
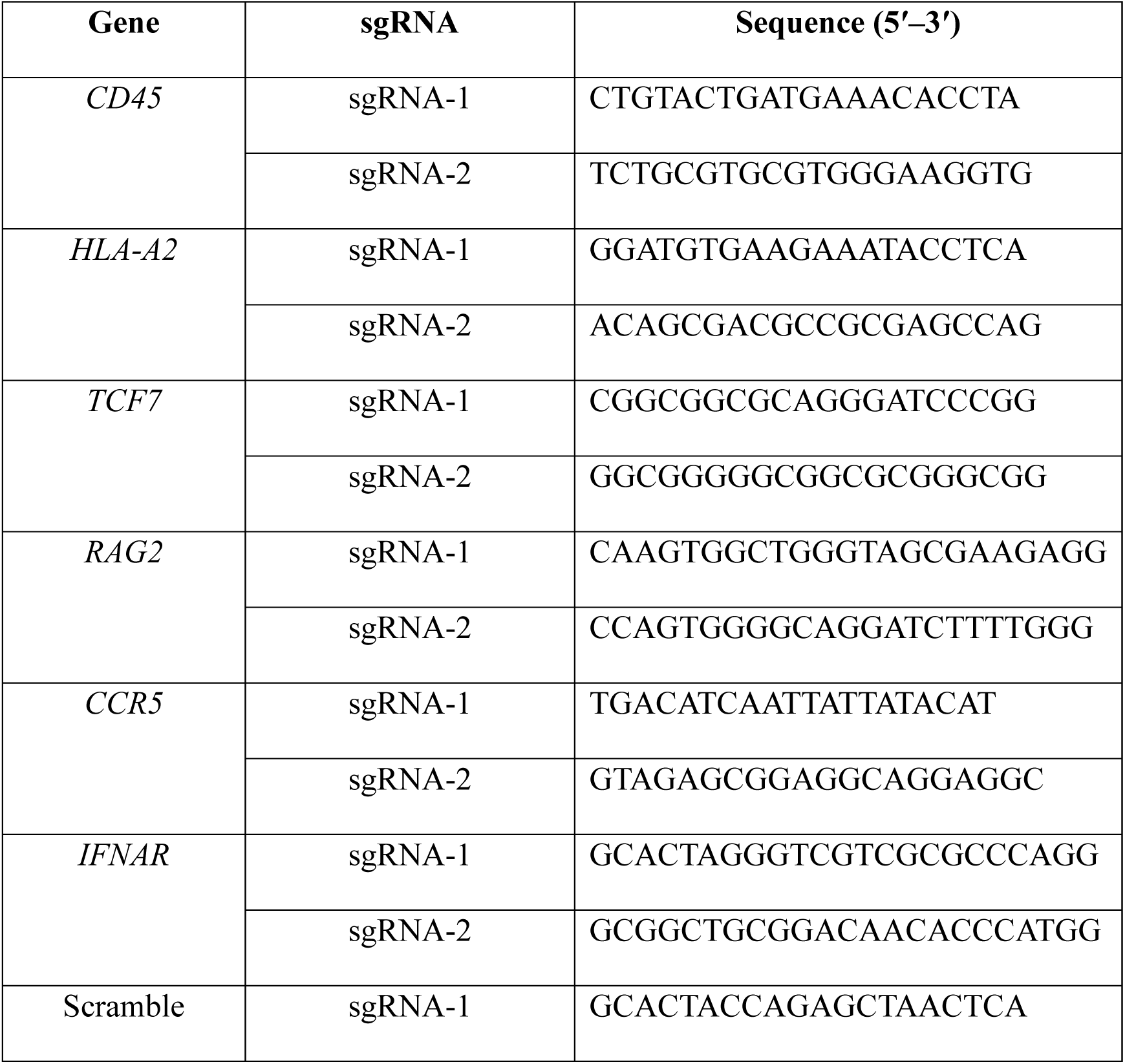
sgRNA sequences.

## References

1 Housden, B. E. et al. Loss-of-function genetic tools for animal models: cross-species and cross- platform differences. Nat Rev Genet 18, 24–40 (2017). 10.1038/nrg.2016.118

2 Rongvaux, A. et al. Human hemato-lymphoid system mice: current use and future potential for medicine. Annu Rev Immunol 31, 635–674 (2013). 10.1146/annurev-immunol-032712-095921

3 Shultz, L. D., Brehm, M. A., Garcia-Martinez, J. V. & Greiner, D. L. Humanized mice for immune system investigation: progress, promise and challenges. Nat Rev Immunol 12, 786–798 (2012). 10.1038/nri3311

4 Sidney, L. E., Branch, M. J., Dunphy, S. E., Dua, H. S. & Hopkinson, A. Concise review: evidence for CD34 as a common marker for diverse progenitors. Stem Cells 32, 1380–1389 (2014). 10.1002/stem.1661

5 Ito, M. et al. NOD/SCID/gamma(c)(null) mouse: an excellent recipient mouse model for engraftment of human cells. Blood 100, 3175–3182 (2002). 10.1182/blood-2001-12-0207

6 Traggiai, E. et al. Development of a human adaptive immune system in cord blood cell- transplanted mice. Science 304, 104–107 (2004). 10.1126/science.1093933

7 Gundry, M. C. et al. Highly Efficient Genome Editing of Murine and Human Hematopoietic Progenitor Cells by CRISPR/Cas9. Cell Rep 17, 1453–1461 (2016). 10.1016/j.celrep.2016.09.092

8 Kuppers, D. A. et al. Gene knock-outs in human CD34+ hematopoietic stem and progenitor cells and in the human immune system of mice. PLoS One 18, e0287052 (2023). 10.1371/journal.pone.0287052

9 Wu, Y. et al. Highly efficient therapeutic gene editing of human hematopoietic stem cells. Nat Med 25, 776–783 (2019). 10.1038/s41591-019-0401-y

10 Wunderlich, M. et al. AML xenograft efficiency is significantly improved in NOD/SCID-IL2RG mice constitutively expressing human SCF, GM-CSF and IL-3. Leukemia 24, 1785–1788 (2010). 10.1038/leu.2010.158

11 Rongvaux, A. et al. Development and function of human innate immune cells in a humanized mouse model. Nat Biotechnol 32, 364–372 (2014). 10.1038/nbt.2858

12 Sungur, C. M. et al. Human NK cells confer protection against HIV-1 infection in humanized mice. J Clin Invest 132 (2022). 10.1172/JCI162694

13 Albanese, M. et al. Rapid, efficient and activation-neutral gene editing of polyclonal primary human resting CD4(+) T cells allows complex functional analyses. Nat Methods 19, 81–89 (2022). 10.1038/s41592-021-01328-8

14 Schumann, K. et al. Generation of knock-in primary human T cells using Cas9 ribonucleoproteins. Proc Natl Acad Sci U S A 112, 10437–10442 (2015). 10.1073/pnas.1512503112

15 Seki, A. & Rutz, S. Optimized RNP transfection for highly efficient CRISPR/Cas9-mediated gene knockout in primary T cells. J Exp Med 215, 985–997 (2018). 10.1084/jem.20171626

16 Schilham, M. W. et al. Critical involvement of Tcf-1 in expansion of thymocytes. J Immunol 161, 3984–3991 (1998).

17 Verbeek, S. et al. An HMG-box-containing T-cell factor required for thymocyte differentiation. Nature 374, 70–74 (1995). 10.1038/374070a0

18 Fugmann, S. D., Lee, A. I., Shockett, P. E., Villey, I. J. & Schatz, D. G. The RAG proteins and V(D)J recombination: complexes, ends, and transposition. Annu Rev Immunol 18, 495–527 (2000). 10.1146/annurev.immunol.18.1.495

19 Berger, E. A., Murphy, P. M. & Farber, J. M. Chemokine receptors as HIV-1 coreceptors: roles in viral entry, tropism, and disease. Annu Rev Immunol 17, 657–700 (1999). 10.1146/annurev.immunol.17.1.657

20 Doms, R. W. Chemokine receptors and HIV entry. AIDS 15 **Suppl 1**, S34–35 (2001). 10.1097/00002030-200102001-00051

21 Margolis, L. & Shattock, R. Selective transmission of CCR5-utilizing HIV-1: the ’gatekeeper’ problem resolved? Nat Rev Microbiol 4, 312–317 (2006). 10.1038/nrmicro1387

22 Shaw, G. M. & Hunter, E. HIV transmission. Cold Spring Harb Perspect Med 2 (2012). 10.1101/cshperspect.a006965

23 Bosinger, S. E. & Utay, N. S. Type I interferon: understanding its role in HIV pathogenesis and therapy. Curr HIV/AIDS Rep 12, 41–53 (2015). 10.1007/s11904-014-0244-6

24 Asmuth, D. M. et al. Safety, tolerability, and mechanisms of antiretroviral activity of pegylated interferon Alfa-2a in HIV-1-monoinfected participants: a phase II clinical trial. J Infect Dis 201, 1686–1696 (2010). 10.1086/652420

25 Lane, H. C. et al. Interferon-alpha in patients with asymptomatic human immunodeficiency virus (HIV) infection. A randomized, placebo-controlled trial. Ann Intern Med 112, 805–811 (1990). 10.7326/0003-4819-112-11-805

26 Rivero, J., Fraga, M., Cancio, I., Cuervo, J. & Lopez-Saura, P. Long-term treatment with recombinant interferon alpha-2b prolongs survival of asymptomatic HIV-infected individuals. Biotherapy 10, 107–113 (1997). 10.1007/BF02678537

27 Sandler, N. G. et al. Type I interferon responses in rhesus macaques prevent SIV infection and slow disease progression. Nature 511, 601–605 (2014). 10.1038/nature13554

28 Hiatt, J. et al. A functional map of HIV-host interactions in primary human T cells. Nat Commun 13, 1752 (2022). 10.1038/s41467-022-29346-w

29 Hultquist, J. F. et al. A Cas9 Ribonucleoprotein Platform for Functional Genetic Studies of HIV-Host Interactions in Primary Human T Cells. Cell Rep 17, 1438–1452 (2016). 10.1016/j.celrep.2016.09.080

30 Carrega, P. et al. CD56(bright)perforin(low) noncytotoxic human NK cells are abundant in both healthy and neoplastic solid tissues and recirculate to secondary lymphoid organs via afferent lymph. J Immunol 192, 3805–3815 (2014). 10.4049/jimmunol.1301889

31 Ferlazzo, G. et al. The abundant NK cells in human secondary lymphoid tissues require activation to express killer cell Ig-like receptors and become cytolytic. J Immunol 172, 1455–1462 (2004). 10.4049/jimmunol.172.3.1455

32 Gautier, E. L. et al. Gene-expression profiles and transcriptional regulatory pathways that underlie the identity and diversity of mouse tissue macrophages. Nat Immunol 13, 1118–1128 (2012). 10.1038/ni.2419

33 Gosselin, D. et al. Environment drives selection and function of enhancers controlling tissue- specific macrophage identities. Cell 159, 1327–1340 (2014). 10.1016/j.cell.2014.11.023

34 Lavin, Y. et al. Tissue-resident macrophage enhancer landscapes are shaped by the local microenvironment. Cell 159, 1312–1326 (2014). 10.1016/j.cell.2014.11.018

35 Masopust, D. & Soerens, A. G. Tissue-Resident T Cells and Other Resident Leukocytes. Annu Rev Immunol 37, 521–546 (2019). 10.1146/annurev-immunol-042617-053214

36 Melsen, J. E., Lugthart, G., Lankester, A. C. & Schilham, M. W. Human Circulating and Tissue- Resident CD56(bright) Natural Killer Cell Populations. Front Immunol 7, 262 (2016). 10.3389/fimmu.2016.00262

37 Ganor, Y. et al. HIV-1 reservoirs in urethral macrophages of patients under suppressive antiretroviral therapy. Nat Microbiol 4, 633–644 (2019). 10.1038/s41564-018-0335-z

38 Ko, A. et al. Macrophages but not Astrocytes Harbor HIV DNA in the Brains of HIV-1-Infected Aviremic Individuals on Suppressive Antiretroviral Therapy. J Neuroimmune Pharmacol 14, 110–119 (2019). 10.1007/s11481-018-9809-2

39 Veenhuis, R. T. et al. Monocyte-derived macrophages contain persistent latent HIV reservoirs. Nat Microbiol 8, 833–844 (2023). 10.1038/s41564-023-01349-3

40 Zalar, A. et al. Macrophage HIV-1 infection in duodenal tissue of patients on long term HAART. Antiviral Res 87, 269–271 (2010). 10.1016/j.antiviral.2010.05.005

41 Pellin, D. et al. A comprehensive single cell transcriptional landscape of human hematopoietic progenitors. Nat Commun 10, 2395 (2019). 10.1038/s41467-019-10291-0

42 Clement, K. et al. CRISPResso2 provides accurate and rapid genome editing sequence analysis. Nat Biotechnol 37, 224–226 (2019). 10.1038/s41587-019-0032-3

